# Positive and negative selective assortment in pairwise interactions

**DOI:** 10.1101/2024.07.18.604169

**Authors:** Segismundo S. Izquierdo, Luis R. Izquierdo, Christoph Hauert

## Abstract

In populations subject to evolutionary processes, the assortment of players with different genes or strategies can have a large impact on players’ payoffs and on the expected evolution of each strategy in the population. Here we consider assortment generated by a process of partner choice known as *selective assortment*. Under selective assortment, players looking for a mate can observe the strategies of a sample of potential mates or co-players, and select one of them to interact with. This selection mechanism can generate positive assortment (if there is a preference for players using the same strategy), or negative assortment. We study the impact of selective assortment in the evolution and in the equilibria of a population, providing results for different games under different evolutionary dynamics (including the replicator dynamics).

## 1 Introduction

The standard model for random encounters of agents in Evolutionary Game Theory (Sandholm, 2010; Weibull, 1995) assumes uniform random matching in large (technically, infinite) well-mixed populations, meaning that any agent is equally likely to meet any other agent. Thus, under uniform random matching, the probability of interacting with an agent who uses strategy *i* (an *i*-player) equals the fraction of *i*-players in the population. Many scholars have pointed out that such situations are probably rare in nature, and argued in favor of studying deviations from the well-mixed model.

A first natural extension of the well-mixed model is to let the probability that two players interact depend on their individual strategies. For instance, if there is *positive assortment*, individuals preferentially interact with individuals of the same type; on the other hand, if there is *negative assortment*, individuals preferentially interact with individuals of a different type. We consider processes in which the average assortment of a type determines its expected payoff, and expected payoffs determine population dynamics. An alternative and more detailed way of departing from the framework of well-mixed populations is to assume that players are embedded on an underlying network. In networks, the probability that a certain individual interacts with other individuals depends on the network configuration (the distribution of strategies over the locations of the network), and local assortment can present considerable fluctuations with respect to average assortment.

Eshel and Cavalli-Sforza (1982) discuss two potential sources of assortment, focusing on positive assortment. The first source is called *structural assortment*, and is associated with situations in which players with different strategies happen to find themselves in different mating environments. This could be due, for instance, to spatial effects: descendants, who share common traits, are usually in close spatial proximity. Local reproduction or local imitation, combined with local interactions, also tend to generate positive assortment.

The second source of assortment is called *selective assortment*, and it assumes that, when looking for a mate or partner, players can meet a (small) number *k* of potential mates, observe their strategy (or some reliable and highly correlated proxy) and select one of those potential mates.

Naturally, both sources of assortment (structural and selective) can take place simultaneously: players may actively select mates in different potential-mate environments.

A reference model for positive assortment is the so-called *two-pool assortative matching process with constant assortativity* α (Bergstrom, 2003, 2013; Eshel & Cavalli-Sforza, 1982), which Eshel and Cavalli-Sforza (1982) interpret as a model of structural assortment. This model assumes that a player interacts:

- with probability α > 0, with a player who uses the same strategy, and
- with probability (1 − α), with a random player from the population.

Equivalently (in terms of expected payoffs), one can suppose that all players in the population are matched in pairs, in a way such that a fraction α of the population is matched assortatively to individuals of their same strategy, and a fraction (1 − α) is matched uniformly at random.

Most models of assortment in the literature on population games (Alger & Weibull, 2010, 2013; Grafen, 1979; Holdahl & Wu, 2023; Iyer & Killingback, 2020; Nax & Rigos, 2016; Newton, 2017) have focused on variations of the two-pool positive assortment process with constant assortativity. As indicated before, this process can be understood as a result of matching or allocating all players into pairs. Jensen and Rigos (2018) provide a general framework for matching rules that allocate all individuals into groups with different compositions (i.e., different frequencies for each strategy), and van Veelen (2011) studies the replicator dynamics in two-strategy games with allocation into groups. Wu (2016) studies two-strategy coordination games in which the index of assortativity is chosen by majority voting.

In this paper we focus on *selective* assortment, which does not assume that all players are matched in pairs. Specifically, we extend Eshel and Cavalli-Sforza‘s (1982) model of selective assortment to allow for more than two strategies and also for negative assortment. Interestingly, to the best of our knowledge, there are no reference models for negative assortment. Probably, one of the reasons lies in the difficulties of extending matching processes with constant assortativity to negative assortment (Jensen & Rigos, 2018). In particular, in appendix A we show that extending the *two-pool assortative matching process with constant assortativity* to model negative assortment can give rise to several undesirable issues, both from a mathematical point of view (discontinuous payoff functions) and in terms of obtaining realistic models.

For the two-strategy case, following a different but related approach, Taylor and Nowak (2006) discuss replicator dynamics with non-uniform interaction rates. These non-uniform interaction rates can also be interpreted in terms of assortment. Interestingly, the phase portraits obtained under selective assortment that we present for the specific case of two strategies and replicator dynamics show some parallels with the phase portraits in Taylor and Nowak (2006). Friedman and Sinervo (2016) present a general framework for assortative interactions based on matching (or encounter) matrices, whose terms can also be interpreted as measures of the frequency with which *i*-players receive payoffs from interactions with *j*-players. Hauert and Miękisz (2018) consider a model in which players who interact together are also more likely to be competitors for reproduction, which leads to deviations from well-mixed populations.

Finally, all phase portraits shown in this paper can be easily replicated with open-source freely available software which performs exact computations of rest points and exact linearization analyses (Izquierdo, Izquierdo, & Hauert, 2024).

## 2 Setting and notation

We consider a population of individuals who may interact with each other in pairs, to play a symmetric two-player game. Each individual has a type or strategy *i* ∈ *S* = {1, 2, …, *n*}. An *i*-player who interacts with a *j*-player obtains a payoff *U*_*ij*_.

Let *x*_*i*_ be the proportion or fraction of type *i* in the population and let ***x*** = (*x*_*i*_)_*i*∈*S*_ be the *population state*: the vector describing the distribution of types in the population. Since *x*_*i*_ ≥ 0 and Σ _*i*∈*S*_ *x*_*i*_ = 1, the population state ***x*** lives in the simplex 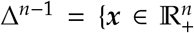: ∑_*i*∈*S*_ *x*_*i*_ = 1}. The monomorphic states in which all players use the same strategy *i* are represented by the unit vectors **e**_*i*_.

At a population state ***x***, each type or strategy *i* is assumed to have an average or expected payoff

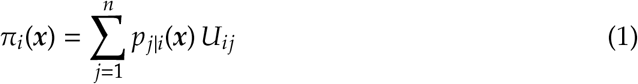

where *p*_*j*|*i*_(***x***) is the conditional probability that an *i*-player interacts with a *j*-player receiving payoff *U*_*ij*_. Thus, in this setting, we allow the probabilities of interaction *p*_*j*|*i*_(***x***) to depend on individual’s *i* type. In this way, we generalize from the standard framework of well-mixed populations where *p*_*j*|*i*_(***x***) = *x*_*j*_ for all *i, j* ∈ *S*. We refer to this last case as *neutral assortment*.

The conditional probabilities *p*_*j*|*i*_(***x***), with 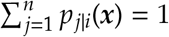, define the assortment of interactions and determine the expected payoff for each strategy type, at any population state ***x***. We assume that these probabilities are defined at every population state ***x*** ∈ Δ^*n*−1^.

Note that the expected payoff π_*i*_(***x***) for strategy *i* (1) is a convex combination of the game payoffs {*U*_*ij*_}, so π_*i*_(***x***) is a value in the range between the minimum and the maximum payoffs for *i*, i.e., π_*i*_(***x***) ∈ [min_*j*_ *U*_*ij*_, max_*j*_ *U*_*ij*_].

### 2.1 Positive and negative assortments

In the literature, the terms *assortment* or *assortative mating* are often used to indicate that individuals interact with their own type with more probability than under random matching. Some authors (see e.g. Iyer and Killingback (2020)) distinguish between positive and negative assortment, depending on whether the conditional interaction probabilities *p*_*i*|*i*_ are greater or lower than under random matching. In the following, we use the term assortment to refer to the set of functions *p*_*j*|*i*_ : Δ^*n*−1^ → [0, 1] that characterize the conditional interaction probabilities (for each strategy pair) at every state. By comparison with the probabilities under random matching, we say that an assortment is:

- Positive if *p*_*i*|*i*_(***x***) ≥ *x*_*i*_ for every *i* ∈ *S* and every state, with strict inequality at least at one state.
- Negative if *p*_*i*|*i*_(***x***) ≤ *x*_*i*_ for every *i* ∈ *S* and every state, with strict inequality at least at one state.
- Neutral if *p*_*i*|*j*_(***x***) = *x*_*i*_ for every *i, j* ∈ *S* and every state.

When referring to an assortment at a specific state ***x***, we say that an assortment at state ***x*** is

- positive if *p*_*i*|*i*_(***x***) > *x*_*i*_ for every *i* ∈ *S*,
- negative if *p*_*i*|*i*_(***x***) < *x*_*i*_ for every *i* ∈ *S*, and
- neutral if *p*_*i*|*j*_(***x***) = *x*_*i*_ for every *i, j* ∈ *S*.

Note that the reference interaction probabilities (those corresponding to random or neutral matching) depend on the state ***x***. In particular, having large values for every same-type interaction probability *p*_*i*|*i*_(***x***) at a population state does not guarantee that there is positive assortment at that state. For instance, if at some state ***x***, *p*_*i*|*i*_(***x***) = 0.9 for every *i* ∈ *S*, but there is a strategy *j* such that *x*_*j*_ > 0.9, then there is no positive assortment. Similarly, having low values for every same-type interaction probability at a state does not guarantee negative assortment at that state. For instance, if at some state ***x***, *p*_*i*|*i*_(***x***) = 0.1 for every *i* ∈ *S*, but there is a strategy *j* such that *x*_*j*_ < 0.1, then there is no negative assortment.

### 2.2 Balanced and boundary-compatible assortments

In many cases it seems natural to assume that if at some population state ***x*** there are no *j*-players (i.e., if *x*_*j*_ = 0), then the conditional probability of meeting a *j*-player at such a state *p*_*j*|*i*_(***x***) must be 0. An assortment that satisfies this condition is said to be boundary-compatible. Specifically, an assortment is *boundary-compatible* if

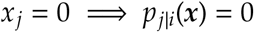

for every state ***x*** ∈ Δ^*n*−1^, and every *i, j* ∈ *S*.

At a monomorphic state **e**_*i*_ (where all players use strategy *i*), the conditional probabilities of a boundary-compatible assortment satisfy *p*_*i*|*j*_(**e**_*i*_) = 1 for every *j* (i.e., if there are only *i*-players, any player entering the population will meet an *i*-player), leading to payoffs π_*j*_(**e**_*i*_) = *U*_*ji*_.

Another interesting property to take into account is *balance*. An assortment may (or may not) imply that, on average, the number of payoff-relevant or accounted *i*-*j* interactions is equal to the number of payoff-relevant *j*-*i* interactions. Specifically, we say that an assortment is *balanced* if:

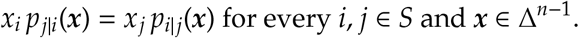

For instance, uniform random matching generates the neutral assortment *p*_*i*|*j*_(***x***) = *x*_*i*_, which is boundary-compatible and balanced: *x*_*i*_ *p*_*j*|*i*_(***x***) = *x*_*i*_ *x*_*j*_ = *x*_*j*_ *p*_*i*|*j*_(***x***). Complete matching (which assumes that every player plays once with every other player) generates the same assortment.

The two-pool process with constant assortativity generates a balanced assortment, but it is not boundary-compatible (α > 0 is a minimum value for the probability *p*_*i*|*i*_ that an *i*-player meets another *i*-player, even at states where there are no *i*-players). Selective assortment, on the other hand, is boundary-compatible, but it is not balanced.

If every time there is an interaction between two players, both players receive a payoff, and all payoffs are treated equally, then the assortment is balanced. This is typically the case if, during a time period, all players in a population are assumed to be matched or grouped in pairs to play the game (whole-population matching). However, there are some reference cases in which assortments are typically non-balanced, as it happens with selective assortment. If players sometimes actively look for an interacting mate and sometimes passively accept a request for interaction, and the relevant payoffs for a player are those obtained when actively looking for an interaction, the associated assortment does not need to be balanced. Similarly, if players of different types have different expected number of interactions per period (e.g., heterogeneous structured populations (Maciejewski, Fu, & Hauert, 2014)) and the relevant payoff is the average payoff per interaction, the associated assortment will typically be non-balanced. In each of these cases some payoffs are treated differently than others, either because of taking averages over a different number of interactions, or because payoffs received by players who do not initiate an interaction are disregarded.

### 2.3 Positive assortment vs positive index of assortativity

For balanced assortments with two types (1 and 2), Bergstrom (2003, 2013) defines the *index of assortativity* at state ***x*** as the difference between the conditional probability of interacting with a type (e.g., type 1) if the player is of that same type (type 1) minus that conditional probability if the player is of the other type (type 2):

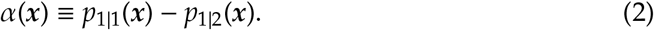

The definition can be extended to non-balanced assortments. It follows from Σ _*j*_ *p*_*j*|*i*_ = 1 that α(***x***) = *p*_1|1_(***x***) − *p*_1|2_(***x***) = *p*_2|2_(***x***) − *p*_2|1_(***x***), and also that α(***x***) = *p*_1|1_(***x***) + *p*_2|2_(***x***) − 1. This last identity leads to the following observation for games with two types:

#### Observation 2.1.

*Positive assortment at* ***x*** *implies positive index of assortativity at* ***x***. *Negative assortment at* ***x*** *implies negative index of assortativity at* ***x***.

For a two-type balanced assortment, it can be shown that *p*_*i*|*i*_(***x***) = *x*_*i*_ + α(***x***)(1 − *x*_*i*_). This implies that, in the special case of balanced assortments, at any interior state ***x*** there is an equivalence between positive (negative) index of assortativity and positive (negative) assortment, as defined in section 2.1.

However, in general, the converse of observation 2.1 is not true: a positive index of assortativity at a state does not guarantee positive assortment at that state, with the corresponding result also for negative assortment. Suppose for instance that at state (*x*_1_, *x*_2_) = (0.5, 0.5) we have *p*_1|1_ = 0.7, *p*_2|1_ = 0.3, *p*_1|2_ = 0.6, *p*_2|2_ = 0.4. At this state we have positive index of assortativity α = 0.7 − 0.6 = 0.4 − 0.3 = 0.1, but there is no positive assortment, because *p*_2|2_ = 0.4 < *x*_2_ = 0.5. In the general case (i.e., when allowing for balanced and non-balanced assortments), having positive assortment at a state is a stronger condition than just having positive index of assortativity at that state.

## 3 Selective assortment

In this section we extend the model of positive selective assortment defined by Eshel and Cavalli-Sforza (1982) for two types, to any number of types and to negative assortment.

Under selective assortment, when players look for a mate they obtain a sample of *k* ≥ 1 random players or potential mates to interact with. The player then chooses as mate:

- Under positive assortment, one of the players who uses her same strategy, if there are any in the sample.
- Under negative assortment, one of the players who uses a different strategy, if there are any in the sample.

If there are no players in the sample with the desired strategies (no players with the same strategy for positive assortment, or no players with a different strategy for negative assortment), then a random mate from the sample is chosen.

For the purpose of the following analysis, the special case *k* = 1, which corresponds to neutral assortment, is included as a reference case in the families of both positive and negative assortment.

### 3.1 Positive Selective Assortment

In this section we derive the assortment probabilities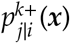 for the model of positive selective assortment with sample size *k* ≥ 1.

The probability that a sample of *k* players has no *i*-players is (1 − *x*_*i*_)^*k*^, so the probability that a sample has at least one *i*-player is 1 − (1 − *x*_*i*_)^*k*^. Considering that every player with a strategy *j ≠ i* is treated equally by an *i*-player who is looking for a mate, it can be shown (see proof in appendix B) that the conditional probabilities 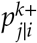 are:

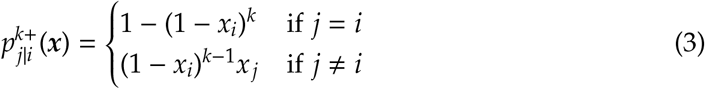

At monomorphic states we have 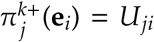. This is so for every sample size *k*, i.e., payoffs at monomorphic states are not affected by the sample size.

At interior states, we have 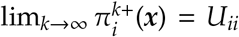. This means that the payoff for every strategy at interior states converges, as *k* → ∞, to the strategy’s same-type-interaction payoff *U*_*ii*_.

### 3.2 Negative Selective Assortment

We now derive the assortment probabilities 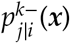 for the model of negative selective assortment with sample size *k* ≥ 1.

The probability that all players in a sample of *k* players are *i*-players is 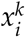. Considering that every player with a strategy *j ≠ i* is treated equally by an *i*-player who is looking for a mate, it can be shown (see proof in appendix B) that the conditional probabilities 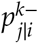 are:

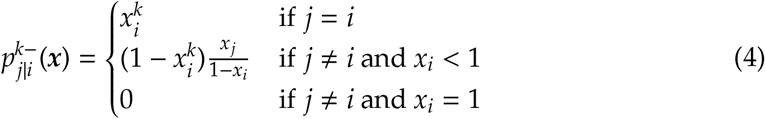

At monomorphic states we have 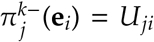. This is so for every sample size *k*, i.e., payoffs at monomorphic states are not affected by the sample size.

At interior states, as *k* → ∞, the probability of same type interactions 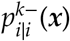 tends to 0, and 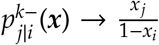 (for *j ≠ i*), i.e., the probability that an *i*-player (when calculating her payoff) interacts with a *j*-player approaches the relative frequency of *j*-players among not-*i*-players.

## 4 Selective assortment in games with two strategies

In this section we analyze the impact of (positive and negative) selective assortment in 2-player 2-strategy symmetric games (henceforth, 2×2 games). We name the strategies *C* for Cooperate and *D* for Defect, and characterize the population state by the fraction of *C*-players *x*_*c*_. Payoffs are *U*_*CC*_ = *R, U*_*DD*_ = *P, U*_*DC*_ = *T*, and *U*_*CD*_ = *S* (see table 1).

**Table 1:** Payoff matrix for a symmetric game with strategies *C* and *D*.

|  |  |  |
| --- | --- | --- |
| | $C$ | $D$ |
| $C$ | $R$ | $S$ |
| $D$ | $T$ | $P$ |

Let us study the different cases, focusing on generic games (i.e., assuming that the four payoffs are different). Without loss of generality, let us assume that coordinating on playing *C* is more efficient than coordinating on playing *D*, i.e., *R* > *P*.

### 4.1 Payoff functions

From (1) and (3), the payoff functions for positive selective assortment are

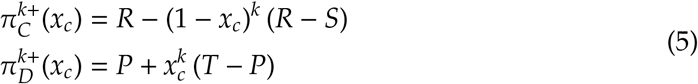

From (1) and (4), the payoff functions for negative selective assortment are

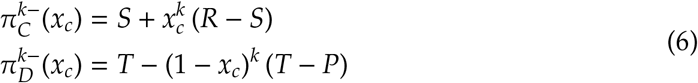

Figure 1 shows some illustrative examples of these functions for a Snowdrift game (*T* > *R* > *S* > *P*).

**Figure 1:**
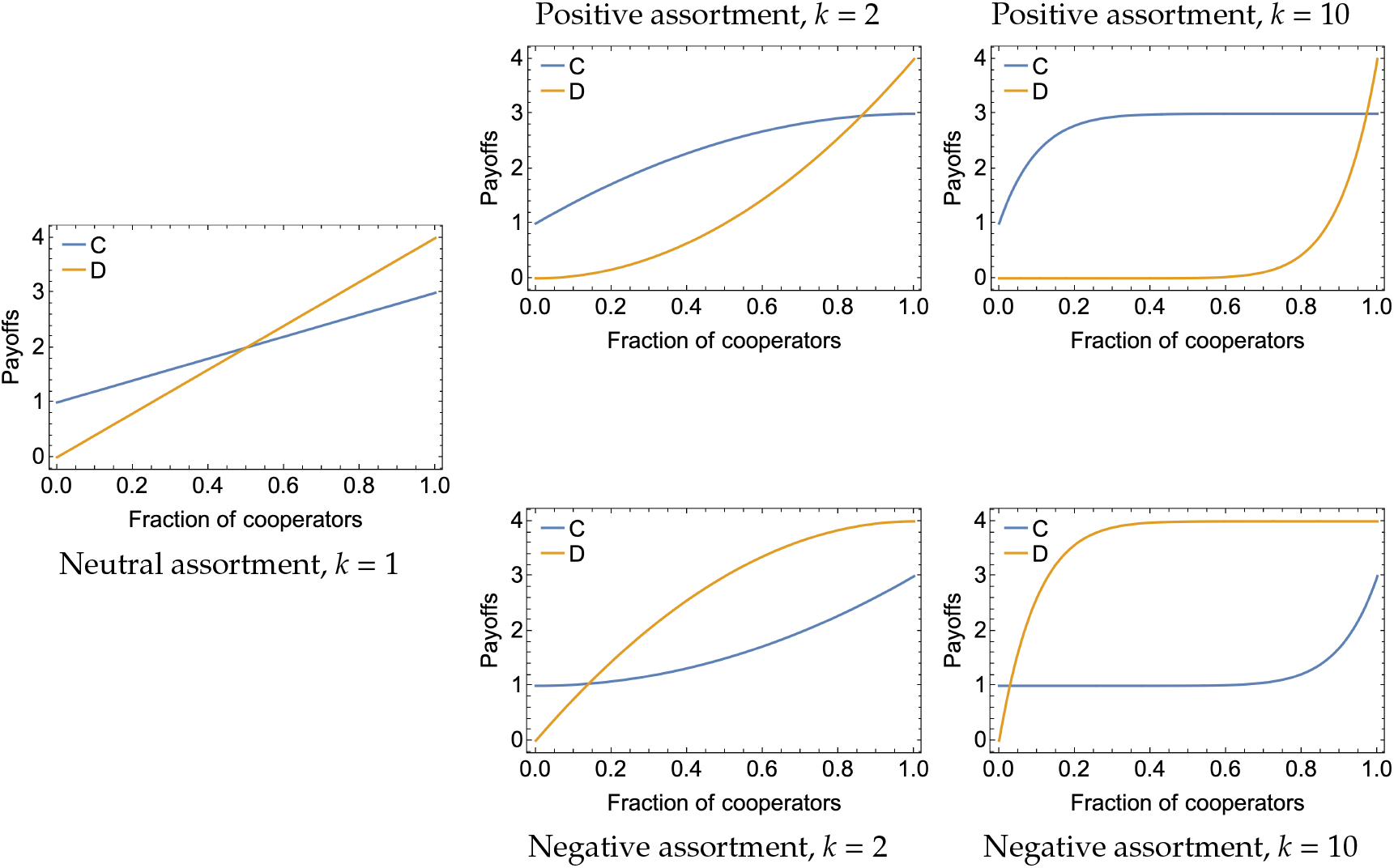
Payoffs for each strategy as a function of the fraction of cooperators in the Snowdrift game with payoffs {*P* = 0, *S* = 1, *R* = 3, *T* = 4} under neutral assortment (*k* = 1), and under positive and negative selective assortment with sample sizes *k* = 2 and *k* = 10. The unique interior ESS moves with *k* in opposite directions depending on whether the assortment is positive or negative.

**Figure 2:**
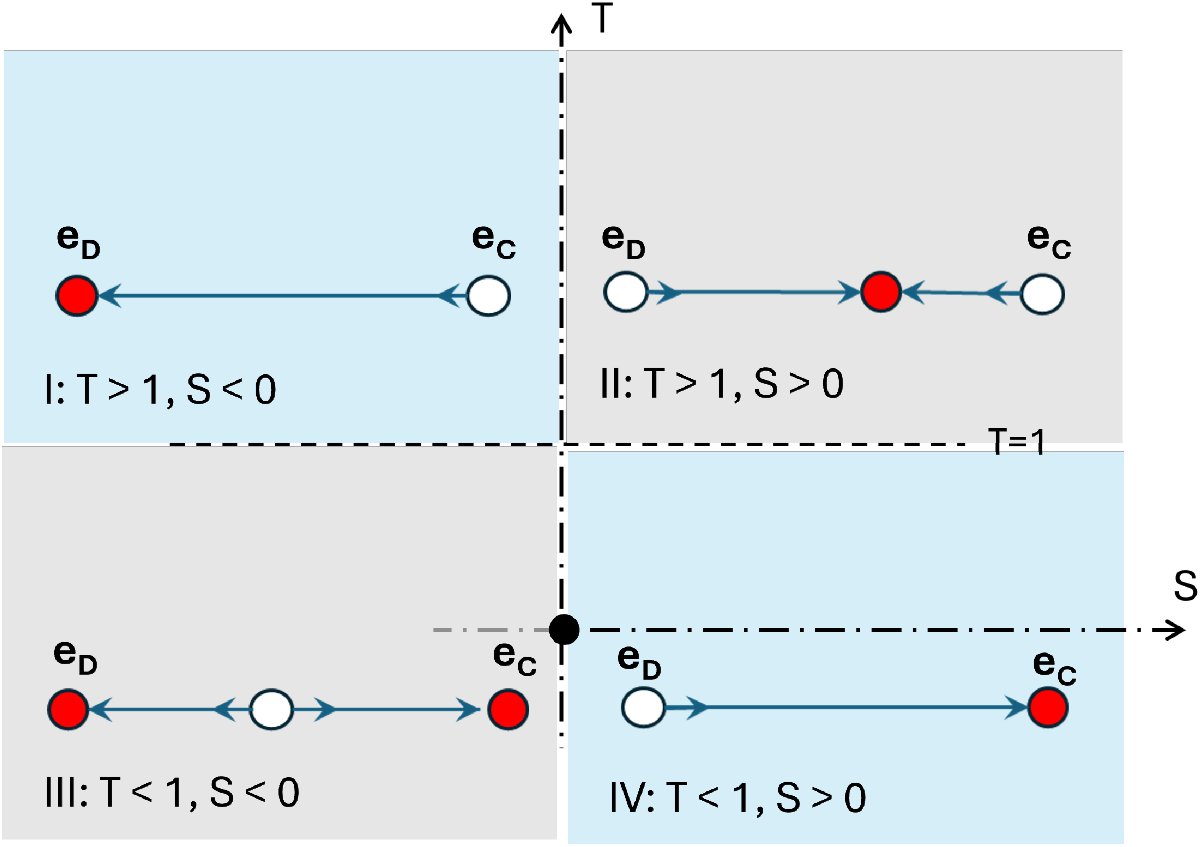
Phase portraits for 2×2 symmetric games in the RD under neutral assortment. Red dots represent evolutionarily stable states (attractors). White dots represent repellors.

It can be seen in the formulas for the payoffs that, at every interior state:

- For positive assortment, as *k* grows, the payoff of each strategy *i* converges to its same-type-payoff *U*_*ii*_ (its main-diagonal payoff in the payoff matrix):

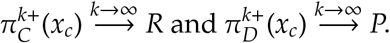
- For negative assortment, as *k* grows, the payoff of each strategy *i* converges to its different-type-payoff *U*_*ij*_ (its anti-diagonal payoff in the payoff matrix):

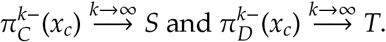

It can also be seen that the payoff functions in (5) and (6) (as functions of *x*_*c*_) are monotonic, because of the monotonicity of 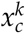 and (1 − *x*_*c*_)^*k*^, with the differences (*R* − *S*) and (*T* − *P*) determining whether the payoff functions are increasing or decreasing, and also determining whether they are convex or concave. For instance, for the Snowdrift game (*R* − *S* > 0, *T* − *P* > 0), 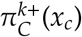 is increasing and concave, while 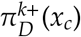 is increasing and convex (see figure 1).

### 4.2 The replicator dynamics

Considering the previous properties of the payoff functions (monotonicity and convexity or concavity), let us study the replicator dynamics

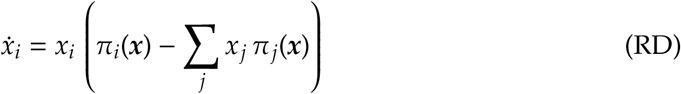

in every 2×2 generic game under selective assortment. Again, assume without loss of generality that *R* > *P*. The phase portrait for the replicator dynamics (RD) does not change if a constant is added to every payoff, or if all payoffs are multiplied by a same positive constant (see (5) and (6)), so for this analysis, without loss of generality, we can assume normalized payoffs *R* = 1 and *P* = 0. (The original payoffs would be normalized by substrating *P* and dividing by *R* − *P*.) We then define the following four regions:

- Region *I*: *T* > 1 and *S* < 0. Example: Prisoner’s Dilemma.
- Region *II*: *T* > 1 and *S* > 0. Example: Snowdrift.
- Region *III*: *T* < 1 and *S* < 0. Example: Stag Hunt.
- Region *IV*: *T* < 1 and *S* > 0. Example: Harmony.

Under neutral assortment, the dynamics in each region are as follows. In region *I, D* is strictly dominant, so **e**_*D*_ attracts all interior trajectories. In region *II*, there is a unique interior evolutionarily stable rest point that attracts all interior trajectories. In region *III*, there is bi-stability of **e**_*C*_ and **e**_*D*_, with an internal unstable rest point separating their basins of attraction. Finally, in region *IV, C* is strictly dominant, so **e**_*C*_ attracts all interior trajectories.

Under positive assortment, a new phase portrait appears in region *I* (Prisoner’s Dilemma) for large enough sample size (see figure 3). This new phase portrait presents an additional attractor (next to **e**_*C*_, for large *k*) and an additional repellor (next to **e**_*D*_, for large *k*). Furthermore, both the level of cooperation and the size of the basin of attraction of this new attractor (where cooperators and defectors coexist) increase as the sample size *k* grows. Note, however, that positive selective assortment cannot stabilize *full* cooperation in the Prisoner’s Dilemma, not even for very large *k*. In region *II*, for large *k*, the interior attractor is close to **e**_*C*_. In region *III*, for large *k*, the interior repellor is close to **e**_*D*_. In every case, for large *k*, the flow in most of the state space points towards **e**_*C*_, which is the most efficient monomorphic state, leading to a stable state (either at **e**_*C*_ or close to it) in which either all or most of the population is cooperating.

**Figure 3:**
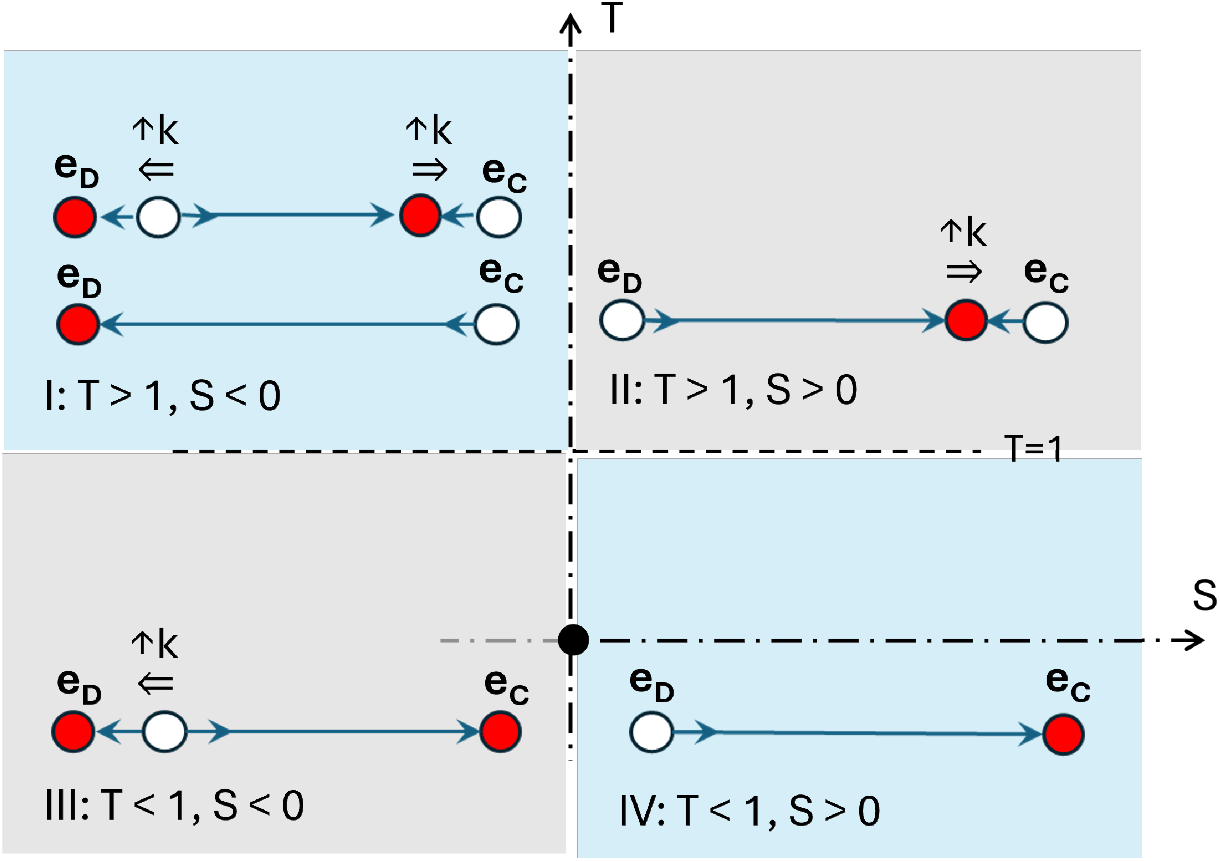
Phase portraits for 2×2 symmetric games in the RD under positive selective assortment. The symbol (↑ *k*) over an interior rest point, with an associated double arrow below, indicates that, for large values of *k*, the interior rest point is close to the corresponding edge.

Under negative assortment, in region *I* (Prisoner’s Dilemma) there is no significant change with respect to neutral assortment (see figure 4). In region *II*, for large *k*, the interior attractor is either close to **e**_*C*_ (if *S* > *T*) or close to **e**_*D*_ (if *T* > *S*). In region *III*, for large *k*, the interior repellor is either close to **e**_*D*_ (if *S* > *T*) or close to **e**_*C*_ (if *T* > *S*). Finally, in region *IV*, a new phase portrait appears for large enough sample size and *T* > *S* (shaded triangle in figure 4). This phase portrait presents an additional attractor (next to **e**_*D*_, for large *k*) and an additional repellor (next to **e**_*C*_, for large *k*). In every case, for large *k*:

**Figure 4:**
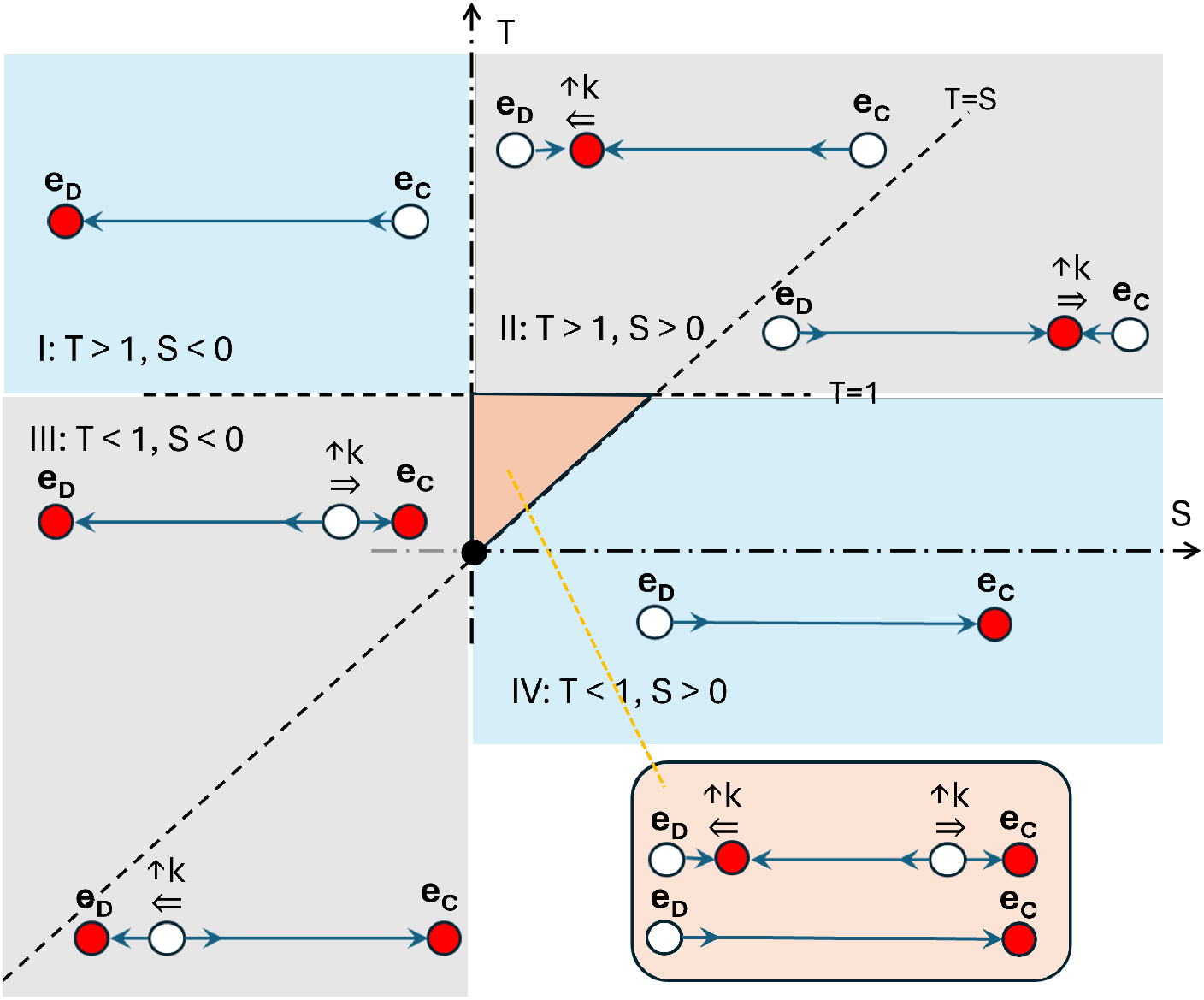
Phase portraits for 2×2 symmetric games in the RD under negative selective assortment. The symbol (↑ *k*) over an interior rest point, with an associated double arrow below, indicates that, for large values of *k*, the interior rest point is close to the corresponding edge.

- If *T* < *S*, the flow in most of the state space points towards **e**_*D*_, leading to a stable state (either at **e**_*D*_ or close to it) in which either all or most of the population is using strategy *D*.
- If *T* < *S*, the flow in most of the state space points towards **e**_*C*_, leading to a stable state (either at **e**_*C*_ or close to it) in which either all or most of the population is using strategy *C*.

## 5 Selective assortment in games with any number of strategies

In this section we consider symmetric two-player games with any number of strategies. We provide results about the equilibria and the attractors, under different evolutionary dynamics, for positive and negative selective assortment.

We will use the concept of a symmetric Nash strategy profile (a profile being a pair of strategies) from classical game theory, and the concept of Nash population state (which is a distribution of strategies in the population) from evolutionary game theory. For completeness, we formally define these concepts below, before indicating their relationship under selective assortment.

### Definition 5.1.

*A strategy profile* (*i, i*) *is a Nash profile if U*_*ji*_ ≤ *U*_*ii*_ *for every j ≠ i. It is a strict Nash profile if the condition holds with strict inequality*.

In the context of population games, considering a set of payoff functions that provide the payoff π_*i*_(***x***) obtained by each strategy at every state, we have the following standard definition of Nash and strict Nash monomorphic states.

### Definition 5.2.

*A monomorphic state* ***e***_*i*_ *is a Nash state if* π_*j*_(***e***_*i*_) ≤ π_*i*_(***e***_*i*_) *for every j ≠ i. It is a strict Nash state if the condition holds with strict inequality*.

### Observation 5.3.

*A monomorphic state* ***e***_*i*_ *is a (strict) Nash state of a game with selective assortment if and only if strategy profile* (*i, i*) *is a (strict) Nash profile of the game*.

With Lipschitz continuous payoff functions (which, given (3) and (4), is the case under selective assortment), strict Nash states are asymptotically stable in the replicator dynamics (Hofbauer & Sigmund, 2003). We consequently have the following result.

### Observation 5.4.

*If* (*i, i*) *is a strict Nash profile, then the monomorphic state* ***e***_*i*_ *is asymptotically stable in the replicator dynamics under selective assortment (either positive or negative, and for every sample size k)*.

Besides the replicator dynamics, observation 5.4 extends to every dynamics for which strict Nash states are asymptotically stable, such as best response dynamics, payoff monotonic imitation dynamics and, more generally, any myopic adjustment dynamics (Hofbauer & Sigmund, 2003).

### Observation 5.5.

*If* (*i, i*) *is not a Nash profile, then the monomorphic state* ***e***_*i*_ *is an unstable rest point of the replicator dynamics under selective assortment (either positive or negative, and for every sample size k)*.

Besides the replicator dynamics, observation 5.5 extends to every dynamics for which non-Nash states are unstable, such as every payoff monotonic imitation dynamics. Furthermore, for many dynamics, such as best response dynamics, only Nash states can be rest points (Sandholm, 2010), so if (*i, i*) is not Nash, then **e**_*i*_ is not even a rest point under such dynamics.

Our next result shows that, for positive assortment and large *k*, there is an attractor of the replicator dynamics close to the most efficient monomorphic state, and most trajectories converge to this attractor.

### Proposition 5.6.

*Assume positive selective assortment. Suppose that there is a unique most-e*ffi*cient monomorphic state* ***e***_*i*_, *i.e*., *U*_*ii*_ > max_*j≠i*_ *U*_*jj*_. *Note that this is always the case in generic games. Then:*

- *At any interior point* ***x***, *for large enough sample size k, strategy i becomes strictly dominant, i.e*,

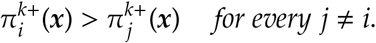
- *Any relative neighborhood O of* ***e***_*i*_ *contains, for large enough k, an attractor of the replicator dynamics. And any trajectory that begins at a state* ***x*** *with x*_*i*_ > 0 *converges, for large enough k, to O. (Note that* ***e***_*i*_ *may not be Nash, and, consequently, it may be unstable.)*

Convergence to a (small) neighborhood of **e**_*i*_ implies that most of the population will adopt strategy *i*, but it does not imply convergence to a rest point.

### Proposition 5.6

extends to every dynamics satisfying

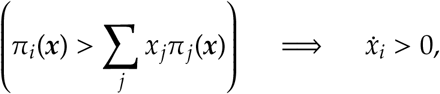

such as aggregate monotonic imitation dynamics (Hofbauer & Sigmund, 2003).

*Example 5.1*. In the Prisoner’s Dilemma (see figure 5), the full-defection state **e**_*D*_ is a strict Nash state, so it is asymptotically stable in the replicator dynamics under every selective assortment. The cooperative state **e**_*C*_ is not Nash, so it is unstable in the replicator dynamics under every selective assortment. However, the most efficient monomorphic state is the cooperative state **e**_*C*_, so, for positive assortment and large enough *k*, there is an attractor close to the cooperative state which attracts most trajectories (see figure 5 for *k* = 10).

**Figure 5:**
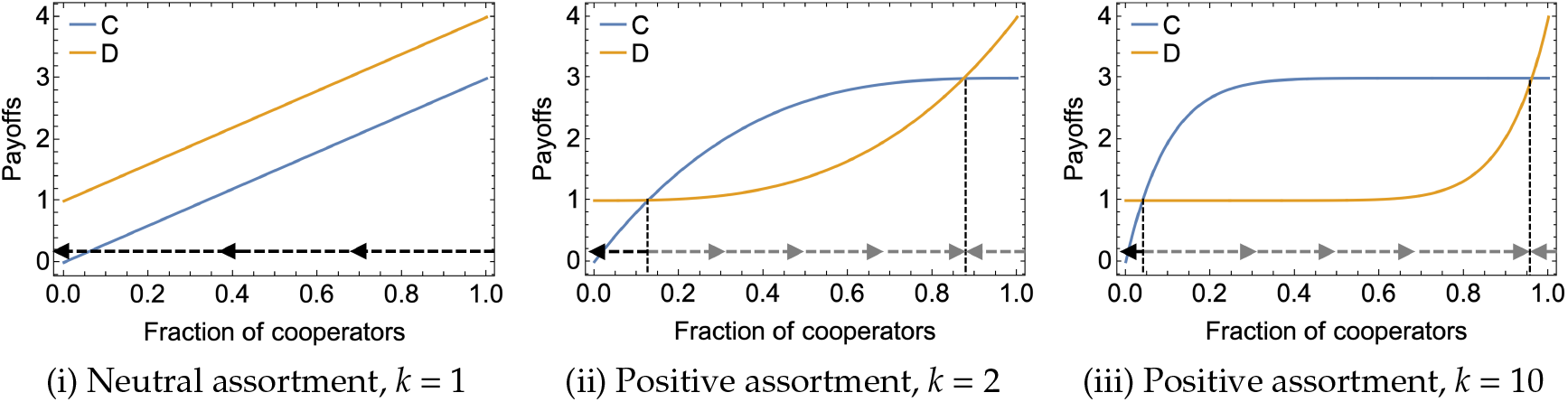
Payoffs for each strategy as a function of the fraction of cooperators in the Prisoner’s Dilemma with payoffs {*S* = 0, *P* = 1, *R* = 3, *T* = 4} under neutral assortment (*k* = 1) and under positive selective assortment with sample sizes *k* = 2 and *k* = 10. The arrows show the phase portrait for the replicator dynamics.

*Example 5.2*. Consider the 1-2-3 coordination game (table 2) with positive selective assortment (figure 6).

**Table 2:**
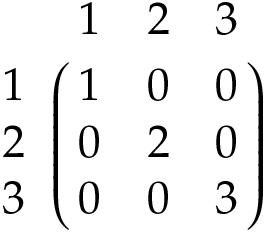
Payoff matrix for the 1-2-3 coordination game.

|  |  |  |  |
|---|---|---|---|
|  | 1 | 2 | 3 |
| 1 | 1 | 0 | 0 |
| 2 | 0 | 2 | 0 |
| 3 | 0 | 0 | 3 |

**Figure 6:**
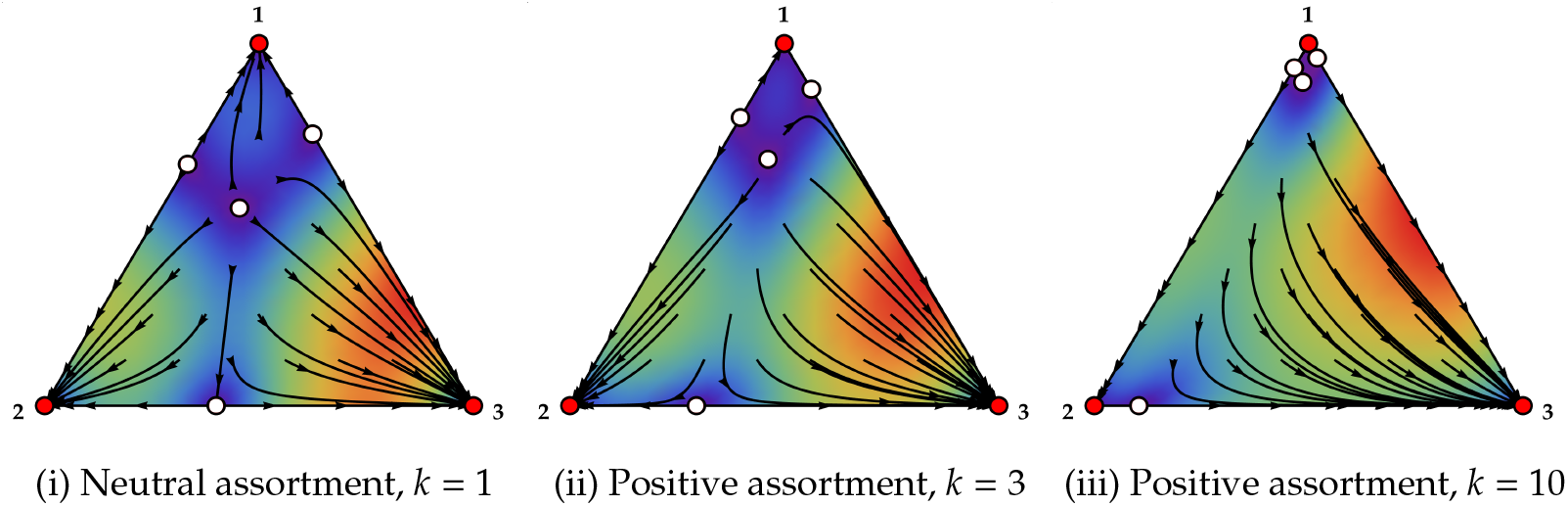
Replicator dynamics for positive selective assortments, for the 123-coordination game with payoff matrix shown in table 2.

The three monomorphic states are strict Nash states, so they are asymptotically stable in the replicator dynamics under (every) selective assortment. The most efficient monomorphic state is **e**_3_. For positive assortment, as *k* grows, it can be seen that most trajectories (all those whose initial value for *x*_3_ is greater than a threshold value that decreases with *k*) converge to the most efficient state **e**_3_.

*Example 5.3*. Consider the Traveler’s dilemma game (table 3), with positive selective assortment (figure 7).

**Table 3:** Payoff matrix for the traveler’s dilemma game with three strategies.

|  | 1 | 2 | 3 |
|---|---|---|---|
| 1 | 2 | 4 | 4 |
| 2 | 0 | 3 | 5 |
| 3 | 0 | 1 | 4 |

**Figure 7:**
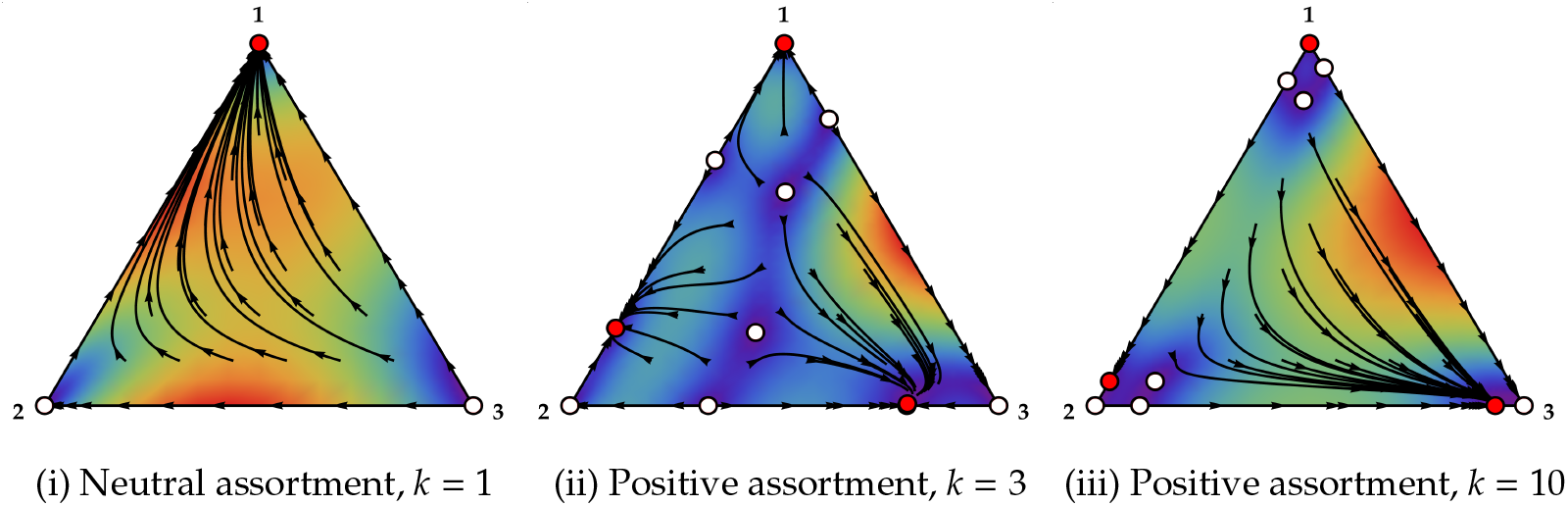
Replicator dynamics for positive selective assortments, for the Traveler’s Dilemma game with the payoff matrix shown in table 3.

Here, the inefficient **e**_1_ is strict Nash, so it is asymptotically stable, while **e**_2_ and **e**_3_ are not Nash, so they are unstable, for every *k*. The least and most efficient monomorphic states are respectively **e**_1_ and **e**_3_. Under neutral assortment, the least efficient state **e**_1_ attracts all interior trajectories. For positive assortment, as *k* grows, it can be seen that most trajectories (all whose initial value for *x*_3_ is greater than some threshold that decreases towards 0 with *k*) converge to an attractor close to the most efficient monomorphic state **e**_3_ (and that attractor gets closer to **e**_3_ as *k* grows).

Our next result, for negative selective assortment and large *k*, provides conditions that guarantee the existence of an attractor close to one of the monomorphic states of a game. It may seem surprising that negative assortment (a preference to interact with other types) leads most players to use the same strategy. However, note that, under selective assortment with large sample size *k*, most interactions in the population take place between players using different strategies even if most players are using the same strategy.

### Proposition 5.7.

*Assume negative selective assortment. If a monomorphic state* ***e***_*i*_ *satisfies* min_*j ≠ i*_ *U*_*ij*_ > max_*j ≠ i*_ *U*_*ji*_, *then any relative neighborhood O of* ***e***_*i*_ *contains, for large enough k, an attractor of the replicator dynamics*.

*If, additionally*, min_*j ≠ i*_ *U*_*ij*_ ≥ max_*j ≠ i,k ≠ j*_ *U*_*jk*_, *then any trajectory that begins at a point* ***x*** *with x*_*i*_ > 0 *converges, for large enough k, to O. (Note that* ***e***_*i*_ *may not be Nash, and, consequently, it may be unstable.)*

Proposition 5.7 extends to any dynamics under which a strictly dominant strategy grows while it is not the only strategy played in the population, i.e., for any dynamics satisfying

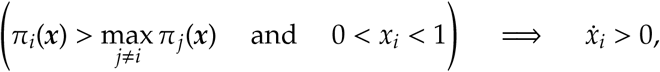

such as best response dynamics.

*Example 5.4*. Consider the game with payoff matrix shown in table 4. Under neutral assortment, strategy 1 is dominated by strategies 2 and 3. Strategy 3 is dominant, and attracts all interior trajectories of the replicator dynamics (see figure 8).

**Table 4:**
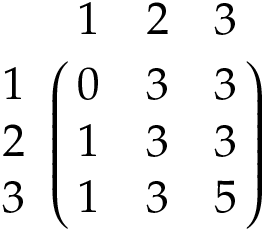
Payoff matrix for the game in example 5.4.

|  | 1 | 2 | 3 |
|---|---|---|---|
| 1 | 0 | 3 | 3 |
| 2 | 1 | 3 | 3 |
| 3 | 1 | 3 | 5 |

**Figure 8:**
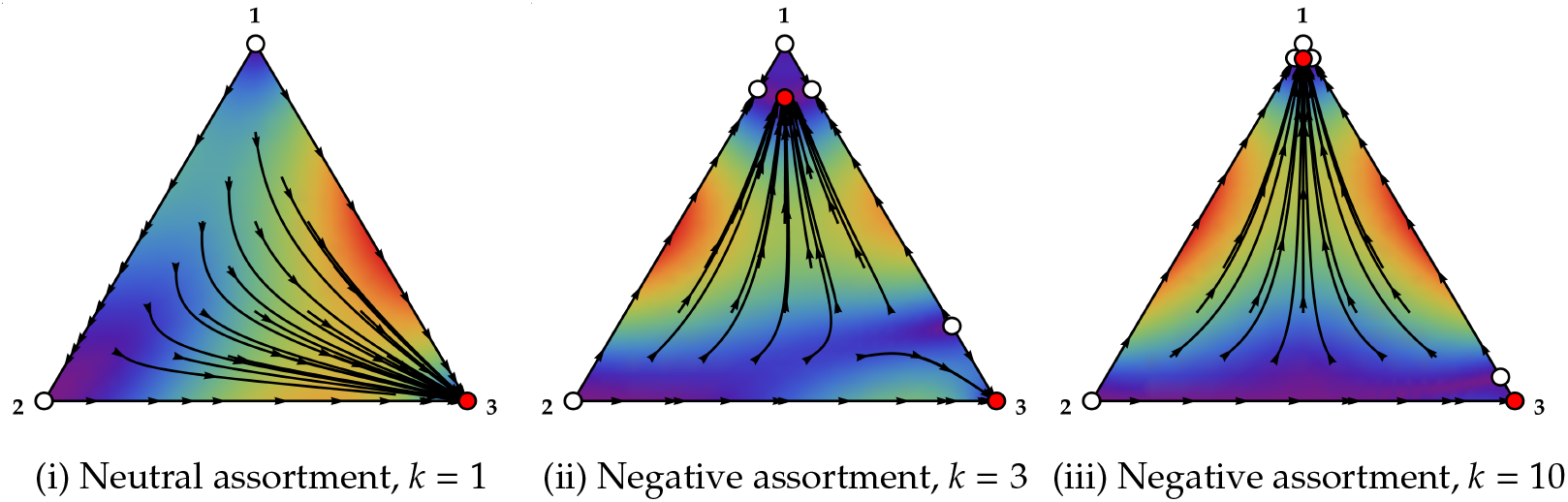
Replicator dynamics for negative selective assortments, for the game in example 5.4.

For strategy 1, after excluding *U*_11_, the minimum row-payoff is greater than the maximum column-payoff (3 > 1). By proposition 5.7, for negative selective assortment and large enough *k*, there is an attractor of the dynamics close to **e**_1_. Furthermore, given that 3 ≥ max_*j ≠* 1,*k ≠ j*_ *U*_*jk*_ = 3, we have that, for ϵ > 0 and large enough *k*, all trajectories with an initial value of *x*_1_ in the range [ϵ, 1 − ϵ] approach this attractor. However, given that **e**_1_ is not a Nash state, **e**_1_ itself is not an attractor.

## 6 Conclusions

In order to study the effects of positive assortment, the *two-pool assortative matching process* (Bergstrom, 2003, 2013; Eshel & Cavalli-Sforza, 1982) can be considered a standard reference model. In sharp contrast, there seems to be no standard reference models to study the effects of negative assortment. Providing reference models for negative assortment is not immediate, because many direct extensions of positive assortment processes (such as extending the two-pool process) can be problematic or unrealistic (see appendix A for details).

In this paper we have analyzed a model of selective assortment for two-player interactions that extends Eshel and Cavalli-Sforza (1982)’s two-strategy positive assortment model to several strategies and, more importantly, to negative assortment. In this way, our contribution fills in the lack of reference models for negative assortment.

Under neutral assortment, in the replicator dynamics with two strategies, the two monomorphic states are rest points and, in generic games, there can be at most one additional interior rest point, which is an attractor or a repellor. Selective assortment (compared to neutral assortment) does not modify the stability of the two monomorphic states, but it can significantly alter the dynamics in the interior of the state space. For instance, in the Prisoner’s Dilemma, if positive selective assortment is sufficiently strong, a new interior attractor appears, where cooperators and defectors coexist. Furthermore, both the level of cooperation and the size of the basin of attraction of this interior attractor increase with the strength of positive assortment. In other games, negative selective assortment can generate a similar effect.

As for results for any number of strategies, we have shown that, under many evolutionary dynamics, positive selective assortment (for sufficiently large sample size *k* and most initial conditions) leads to most of the population playing the most efficient strategy, i.e. the strategy with the greatest same-type payoff *U*_*ii*_. For negative assortment, we have also identified strategies that, under many evolutionary dynamics, are adopted by the majority of the population.

## A Alternative models for negative assortment

Somewhat surprisingly, defining the negative-assortment equivalent of a process that generates positive assortment is not generally straightforward or even possible. In this appendix, we present potential extensions of the *two-pool positive assortment model* (Eshel & Cavalli-Sforza, 1982) to include negative assortment, and we discuss the issues that each of them presents. We start by presenting three possible characterizations of the *two-pool positive assortment model*. Each of these characterizations leads to a different model of negative assortment, which we formally define and discuss.

### A.1 Three characterizations of the two-pool positive assortment model

The following characterizations can be used to define the *two-pool positive assortment model*: [C-I] Players compute their payoffs by interacting:

- with probability α > 0, with a player using their same strategy, and
- with probability (1 − α), with a random player.

[C-II] The assortment is balanced and all *assortativity factors* α_*ij*_(***x***) are equal to a positive constant α, i.e. *p*_*i*|*i*_(***x***) − *p*_*i*|*j*_(***x***) = α for every *i* and *j ≠ i*.

[C-III] At interior states, a representative fraction α of the population is matched in pairs in a way such that the number of same-strategy pairs is maximized, and the remaining fraction (1 − α) of the population is randomly matched.

We now summarize the main properties of the *two-pool positive assortment model* (regardless of how it is characterized). Probabilities *p*_*j*|*i*_(***x***) in this assortment are:

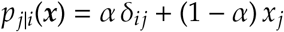

where δ_*ij*_ is the Kronecker delta (δ_*ii*_ = 1, and δ_*ij*_ = 0 if *i ≠ j*).

The payoffs that result from this assortment are linear in ***x*** and are equivalent to the pay-offs obtained under uniform random matching using the modified game payoffs:

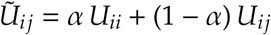

And finally, this assortment is balanced and positive, but not boundary compatible since *p*_*i*|*i*_(***x***) ≥ α at every state. This implies that even if there are no *i*-players in a population (*x*_*i*_ = 0), a potential invader using strategy *i* is assumed to be able to interact with another *i*-player with probability at least α (see Bergstrom (2013) for an alternative model). This is probably the main drawback of this model of positive assortment.

The following subsections present and discuss potential ways to model negative assortment, taking each of the three characterizations of the *two-pool positive assortment model* as a starting point.

### A.2 Extension from [C-I]. Proportional negative assortment

A natural way to model negative assortment in the spirit of characterization [C-I] would be the following:

**Extension from [C-I]**. Players compute their payoffs by interacting:

- with probability α > 0, with a player using a different strategy, and
- with probability (1 − α), with a random player.

In contrast with the case of positive assortment, here we must also specify the probability of selecting each of the different strategies. A natural way of doing this is proportional to their frequencies. In that case, the probabilities *p*_*j*|*i*_(***x***) are:

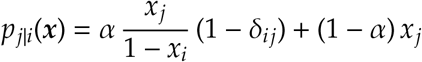

Probabilities *p*_*j*|*i*_(***x***) would still need to be defined at monomorphic states where *x*_*i*_ = 1, but the main issue with this model is that, in generic games with more than two strategies, payoffs are necessarily discontinuous at monomorphic states for any α > 0. In other words, payoffs at monomorphic states cannot be defined in a continuous way. This is so because lim_*ϵ*→0_ π_1_(1 − *ϵ, ϵ*, 0, …, 0) = α*U*_12_ + (1 − α) *U*_11_ and lim_*ϵ*→0_ π_1_(1 − *ϵ*, 0, *ϵ*, 0, …, 0) = α*U*_13_ + (1 − α) *U*_11_.

### A.3 Extensions from [C-II]. Assortment with constant negative assortativity

A natural way to model negative assortment in the spirit of characterization [C-II] would be the following:

**Extension from [C-II]**. The assortment is balanced and all *assortativity factors* α_*ij*_(***x***) are equal to a negative constant −α, i.e. *p*_*i*|*i*_(***x***) − *p*_*i*|*j*_(***x***) = −α for every *i* and *j ≠ i*.

It turns out that there are no balanced assortments with negative constant index of assortativity or, more generally, with negative constant assortativity factors. Jensen and Rigos (2018) prove this fact for the two-strategy case, but the statement also applies to more than two strategies. To see this, consider a state 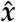 with 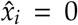 and 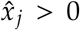. The balancing condition implies 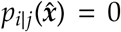, so the corresponding assortativity factor is 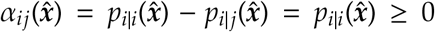. Thus, the assortativity factors of a balanced assortment cannot be a negative constant.

Having seen that a balanced assortment with constant negative assortativity cannot exist, now we explore whether a non-balanced assortment with constant negative assortativity may exist.^1^ The only condition we impose is:

#### Second extension from [C-II]

The assortment is such that all *assortativity factors* α_*ij*_(***x***) are equal to a negative constant −α, i.e. *p*_*i*|*i*_(***x***) − *p*_*i*|*j*_(***x***) = −α for every *i* and *j ≠ i*.

Note that, for *j ≠ i*, this extension implies *p*_*i*|*j*_(***x***) = α + *p*_*i*|*i*_(***x***) > 0. As a consequence, such assortments may not be very realistic because they imply that, in a population with no *i*-players, any not-*i*-player can interact with an *i*-player with strictly positive probability.

### A.4 Extension from [C-III]. Matching with maximum number of different-strategy pairs

A natural way to model negative assortment in the spirit of characterization [C-III] would be the following:

#### Extension from [C-III]

At interior states, a representative fraction α of the population is matched in pairs in a way such that the number of different-strategy pairs is maximized, and the remaining fraction (1 − α) of the population is randomly matched.

The two-strategy case is not problematic. The payoffs obtained when maximizing the number of different-strategy pairs are:

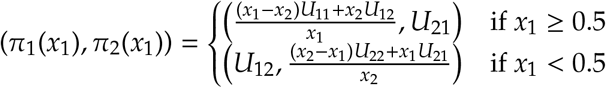

For more than three strategies, problems start to appear. A first issue to be addressed is that, at some population states, there are different matchings in pairs that maximize the number of different-strategy pairs, and these different matching mechanisms can lead to different payoffs (for instance, any matching in pairs of four players who use four different strategies maximizes the number of different-strategy pairs). In any case, for more than two strategies, and considering generic games, this assortment necessarily leads to discontinuous payoff functions, as our next proposition shows.

**Proposition A.1**. *In populations with more than two strategies, and considering generic games, any matching mechanism that maximizes the number of different-strategy pairs leads to discontinuous payoff functions*.

*Proof*. Any matching that maximizes the number of different-strategy pairs satisfies:

- *p*_3|1_ = 1 at states 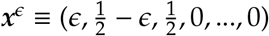, for 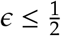. This implies π_1_(***x***^*ϵ*^) = *U*_13_.
- *p*_2|1_ = 1 at states 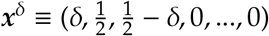, for 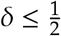. This implies π_1_(***x***^δ^) = *U*_12_.

Let 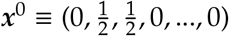. We have lim_*ϵ*→0_ ***x***^*ϵ*^ = ***x***^0^ and lim_*δ*→0_ ***x***^*δ*^ = ***x***^0^, but lim_*ϵ*→0_ π_1_(***x***^*ϵ*^) = *U*_13_ and lim_*δ*→0_ π_1_(***x***^*δ*^) = *U*_12_ * *U*_13_ (because the game is assumed to be generic), so π_1_(***x***) cannot be continuous at *x*^0^. □

## B Proofs

*Proof of equation* (3). We know that 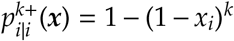 and that the process is such that, at state ***x***, every player with a strategy that is not *i* has the same probability of being chosen by an *i*-player. Consequently, 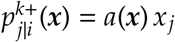 for some value *a*(***x***), so

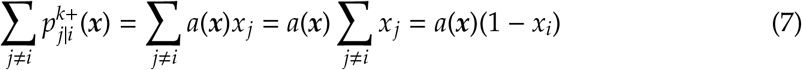

From 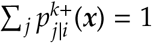 we also have

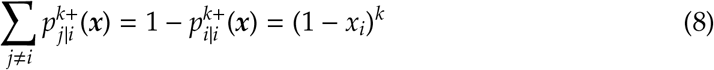

Combining (7) and (8) we find 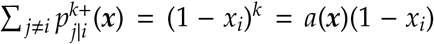 leading to *a*(***x***) = (1 − *x*_*i*_)^*k*−1^. Consequently, 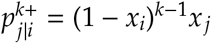 for *j ≠ i*.

*Proof of equation* (4). We know that 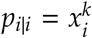 and that the process is such that, at state ***x***, every player with a strategy that is not *i* has the same probability of being chosen by an *i*-player. Consequently, 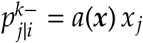 for some value *a*(***x***), and

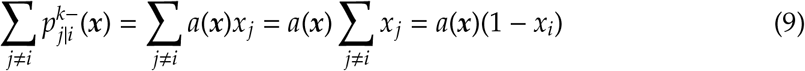

From 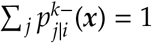, we also have

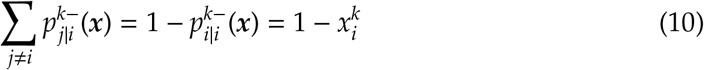

Combining (9) and (10) we find 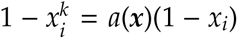, leading to 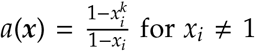 for *x*_*i*_*≠* 1. Therefore, 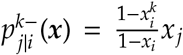 for *j ≠ i* and *x*_*i*_*≠*1. For *x*_*i*_ = 1 we have 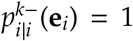 and, consequently, 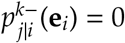 for *j ≠i*. □

*Proof of observation 5.3*. 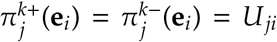 for every *i, j* ∈ *S* and for every *k*, from which the result follows. □

*Proof of proposition 5.6*. From (1) and (3) we have

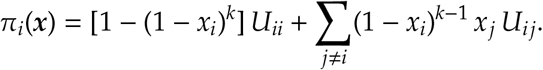

It follows that, at any interior point ***x***, lim_*k*→∞_ π_*i*_(***x***) = *U*_*ii*_, and, if **e**_*i*_ is the most-efficient monomorphic state and *j ≠ i*, lim_*k*→∞_ π_*j*_(***x***) = *U*_*jj*_ < *U*_*ii*_, proving the first part of proposition 5.6.

For the second part, let 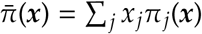 be the average payoff at state ***x***, let *D*_1_ ≡ *U*_*ii*_ = max_*j*_ *U*_*jj*_ and let *D*_2_ ≡ max_*j*\**i*_ *U*_*jj*_ < *D*_1_. We will prove that, for every ***x*** with *x*_*i*_ > 0,

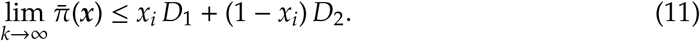

Then, for 0 < *x*_*i*_ < 1 (given that *D*_2_ < *D*_1_),

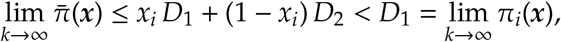

from which the result follows (for *ϵ* > 0, large enough *k* and *x*_*i*_ ∈ [*ϵ*, 1 − *ϵ*], we have 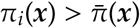, which implies 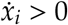 for *x*_*i*_ ∈ [*ϵ*, 1 − *ϵ*]).

It only remains to show (11). In the following bound, we use the value *M* ≡ max_*j≠i,k≠j*_ *U*_*jk*_.

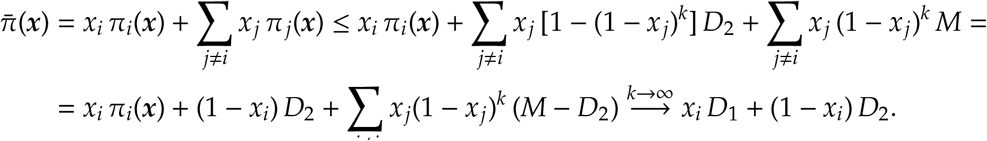

*Proof of proposition 5.7*. From (1) and (4) we have that, for *x*_*i*_ < 1,

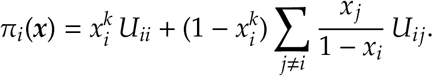

Let *i* be the strategy satisfying the condition min_*j≠i*_ *U*_*ij*_ > max_*j≠i*_ *U*_*ji*_. We use the auxiliary variables *B*_1_ ≡ min_*j≠i*_ *U*_*ij*_, *B*_2_ ≡ max_*j≠i*_ *U*_*ji*_ (so, by hypothesis, *B*_1_ > *B*_2_) and *M* ≡ max_*j≠i,k≠j*_ *U*_*jk*_. At any point ***x*** with *x*_*i*_ < 1 we have

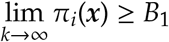

At any point ***x*** with *x*_*i*_ > 0, we have that, for *j ≠ i*,

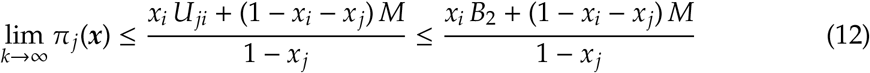

The upper bound in (12) is a convex combination of *B*_2_ and *M*. If *M* ≤ *B*_1_ (and considering that *B*_2_ < *B*_1_), then for every ***x*** with 0 < *x*_*i*_ < 1 we have lim_*k*→∞_ π_*i*_(***x***) > lim_*k*→∞_ π_*j*_(***x***). This implies that, for large enough *k*, strategy *i* is strictly dominant for *x*_*i*_ ∈ [*ϵ*, 1 − *ϵ*] (fixing first *ϵ* > 0, and then taking a large enough *k*), proving the result (for *M* ≤ *B*_1_).

If *M* > *B*_1_, let γ > 0 be a positive constant. We have from (12) that, for *x*_*i*_ > γ,

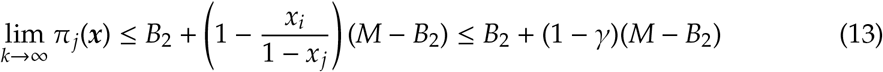

Solving for γ_0_ in

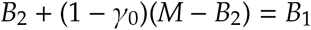

we find 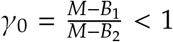, so for 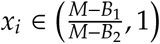, we have lim_*k*→∞_ π_*i*_(***x***) > lim_*k*→∞_ π_*j*_(***x***). We consequently have compact regions 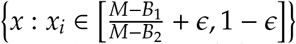, for small enough *ϵ* > 0, in which strategy *i* is strictly dominant for large enough *k*, proving the result. □

## Footnotes

1 For the two-strategy case, Friedman and Sinervo (2016) discuss some balanced assortments with (non-constant) negative assortativity.

